# Seroprevalence of *Toxoplasma gondii* among sylvatic rodents in Poland

**DOI:** 10.1101/2021.01.27.428493

**Authors:** Maciej Grzybek, Daniela Antolová, Katarzyna Tołkacz, Mohammed Alsarraf, Jolanta Behnke-Borowczyk, Joanna Nowicka, Jerzy Paleolog, Beata Biernat, Jerzy M. Behnke, Anna Bajer

## Abstract

*Toxoplasma gondii* is a significant pathogen affecting humans and animals. We conducted seromonitoring for *T. gondii* in four sylvatic rodent species in Poland. We report an overall seroprevalence of 5.5% (3.6% for *Myodes glareolus* and 20% for other vole species). Seroprevalence in bank voles varied significantly between host age and sex.

## INTRODUCTION

There is currently considerable interest in understanding not only the transmission of pathogens but also the range of different variables that influence infection dynamics. Among these, both extrinsic factors (e.g. geographic location) and intrinsic factors, (e.g. host sex) are known to play varying but crucial roles in the exposure of hosts and their susceptibility to infection (1,2). Analyzing pathogen dynamics in their wildlife reservoirs is essential in providing a good understanding of their epidemiology and facilitating informed decisions on appropriate measures for their control. Wild rodents pose a particular threat for human communities because they constitute the most abundant and diversified group of all living mammals (3). Although some zoonotic parasites are not transmissible directly from rodents to humans, or pose just a low risk of direct transmission, predation of rodents by carnivore pets may cause infections in these animals, leading to subsequent contamination of households and other human-related environments (4).

*T. gondii* is an intracellular Apicomplexan parasite with a broad range of intermediate hosts, including humans and rodents (5). The parasite is present first in the tachyzoite stage and later as bradyzoites, also called tissue cysts (6). These can pass from host to host via the food chain and across the placenta resulting in congenitally acquired infections. Rodents are considered to be reservoirs of infection for their predators that include cats, pigs and dogs. Felid species are the definitive hosts of *T. gondii* and the only hosts that can shed *T. gondii* oocysts into the environment

In 2017, 194 confirmed human congenital toxoplasmosis cases were reported from 22 EU countries. France accounted for 79% of all cases. The number of congenital infections per 100 000 newborns was 5.3 in the EU/EEA. The highest incidence was reported in France (19.9), followed by Slovenia (9.98), Poland (4.48) and Bulgaria (3.13) (7). In 2018 only 25 cases of congenital toxoplasmosis were reported in Poland, and in 2019 even fewer with only 14 confirmed cases. (8).

We conducted a multi-site, long-term study on *T. gondii* in northeastern Poland. Our objectives were to monitor seroprevalence of *T. gondii* in the four abundant vole species found in the region (*Myodes glareolus, Microtus arvalis, Microtus agrestis, Alexandromys oeconomus*) and to assess variation in seroprevalence attributable to both intrinsic and extrinsic factors that were quantified.

## MATERIAL AND METHODS

Our study sites are located in the Mazury Lake District region in north-eastern Poland. The sites, methods used for trapping rodents, and for sampling and processing trapped animals have all been thoroughly described (9–11). Bank voles were sampled in 2002, 2006 and 2010 in three local but disparate forest sites. Other species of voles were sampled only in 2013 from fallow meadows close to one of the forest sites. A bespoke ELISA test was used to detect antibodies against *T. gondii* using commercially available *T. gondii* antigen (CD Creative Diagnostics, New York, USA) prepared from *T. gondii* RH strain (12). The statistical approach has been documented comprehensively in our earlier publications (1,13). Prevalence values are given with 95% confidence limits in square brackets. All of the procedures were conducted with the approval of the First Warsaw Local Ethics Committee for Animal Experimentation in Poland.

## RESULTS

We examined 577 rodent individuals, and found *T.gondii* antibodies in the sera of all four rodent species, with an overall seroprevalence of 5.5% (Table 1). There was a significant difference in seroprevalence between vole species (*χ* ^2^_3_=22.58; *P*=0.001) with *Microtus* and *Alexandromys* spp. showing 5.6-fold higher seroprevalence (20% [12-30.9]) than *M. glarolus* (3.6% [2.6-4.9]). In a log-linear model restricted to *Microtus* and *Alexandromys* spp. there was no difference between these three species (*χ* ^2^_2_=0.8, *P*=0.7), nor between the sexes (*χ* ^2^_1_=0.37; *P*=0.55) or age classes *χ* ^2^_2_=4.67; *P*=0.097), although the latter was close to significance and reflected the peak seroprevalence in age class 2 (35.7 [15.28-62.89]) in comparison to zero seroprevalence among the youngest voles and 18.0% [9.46-30.87] among the oldest (Figure 1). We therefore confined further analyses to bank voles *(M. glareolus*).

**Figure 1.**
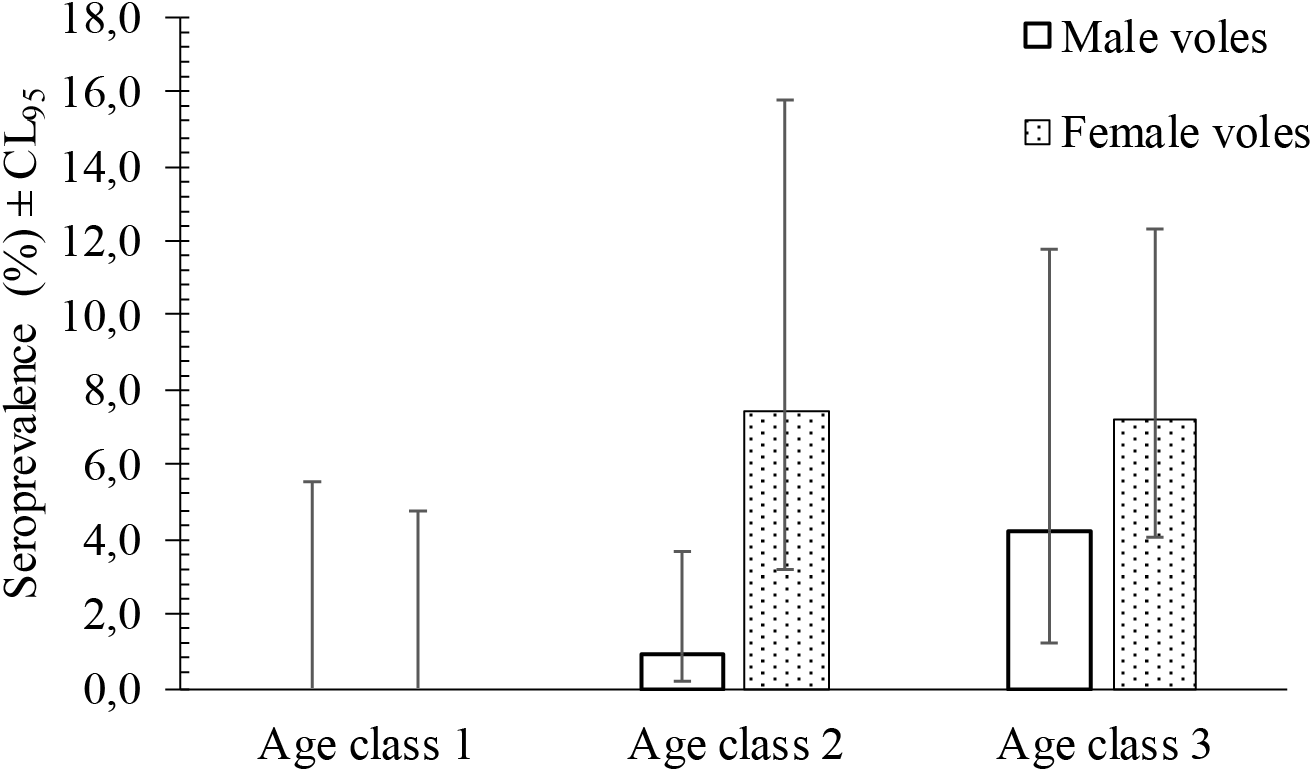
Seroprevalence of *T. gondii* in bank voles in Poland by host sex and by host age class (class 1—immature juvenile voles; class 2—young adult voles; and class 3—breeding older animals). Error bars indicate 95% CL.

**Table 1.**
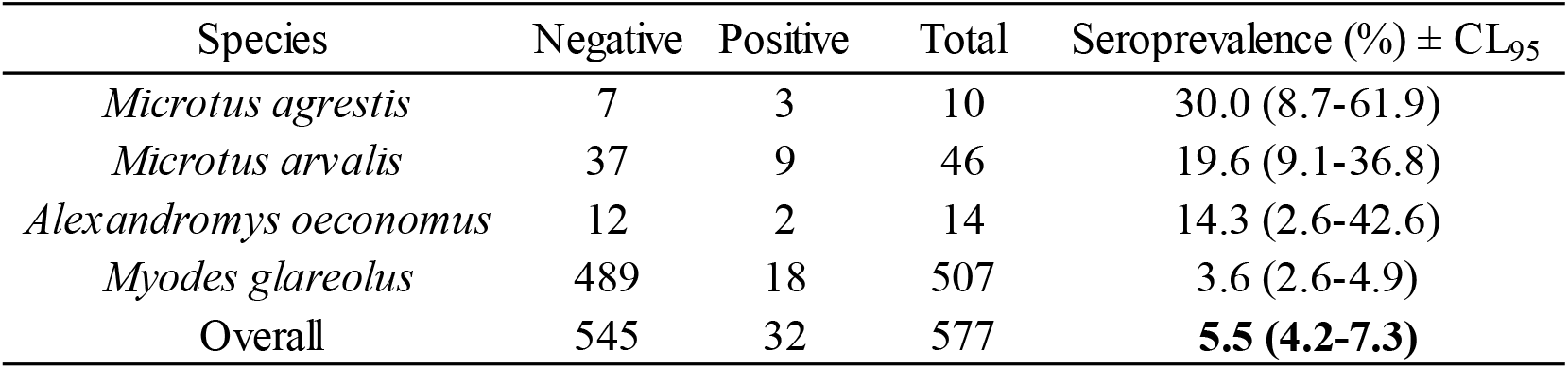
Seroprevalence of *T. gondii* in four rodent species in Poland. Seroprevalence is given in percentages with 95% CL in brackets.

In a log-linear model confined to bank voles, and with year of sampling (3 years) and site (3 sites) taken into account, *T.gondii* seroprevalence differed significantly between the sexes of *M. glareolus* (*χ*^2^_1_=4.34; *P*=0.037) and was 3.5 times higher for female compared with male bank voles (Figure 1). *T. gondii* seroprevalence increased significantly with host age (*χ* ^2^_2_=11.57; *P*=0.003), peaking in the oldest bank voles (6.1% [2.77-12.47]) but was lower in bank voles from age class 2 (mostly young adults; 3.5% [1.22-9.50]). No immature bank voles were found to be seropositive. The data in Figure 1 also indicate that seroprevalence increased with host age faster among female bank voles compared with males, although the interaction between age and sex was not significant (*χ* ^2^_2_=0.28; *P*=0.87).

## DISCUSSION

Our results confirm that both *M. glareolus* and *Microtus/Alexandromys* spp. have contact with *T. gondii* and become infected. Therefore, they may potentially play a role as reservoirs of this parasite in the sylvatic environment. According to the meta-analysis carried out by Galeh and colleagues, the overall seroprevalence of *anti-Toxoplasma* IgG antibodies in rodents in Europe is 1% (14). Here, we report an overall seroprevalence of *T. gondii* in voles of 5.5%, with *Microtus* and *Alexandromys* spp. showing a considerably higher seroprevalence at 20%. Although the reason for this discrepancy between the rodent genera is not clear, we have noted that cats from local farms are often observed in the meadows where we trapped *Microtus/Alexandromys* spp., but seldom enter the forests to which bank voles are confined. This is in line with the results obtained with radio-collared farm cats in Switzerland, which kept a short distance from the house and preferred meadows over forests (15).

We found that female bank voles were 3.5 times more likely to be infected with *T. gondii* than males, and in this respect, our data contrast with reports in the literature in which *T. gondii* seropositivity in male rodents has been recorded generally to be higher than in females (14). It was not unexpected to find in our study that older animals were more likely to have experienced infection than juveniles, since the current work was based on the presence/absence of specific antibody against *T. gondii*, reflecting the history of previous infections and not necessarily a current infection. Older animals will have had a longer time frame over which to encounter the infective stages of *T. gondii*, and hence greater likelihood of infection than younger individuals.

Transmission of the parasite from wildlife to human habitats may occur especially in rural areas where cats escape human settlements and forage on wild rodents. Further studies are necessary therefore, to reveal comprehensively the status of toxoplasmosis in wildlife and to assess the risk of infection for local inhabitants, as well as for visitors to the region.

## ACKNOWLEDGEMENTS

We thank Dr Ewa Mierzejewska for her help in rodent trapping and the fieldwork. We also thank the University of Nottingham, University of Warsaw, Institute of Parasitology, Slovak Academy of Sciences and the Medical University of Gdańsk for financial support. JMB was supported by the Royal Society, the British Ecological Society and the Grabowski Fund. AB was supported by the Polish State Committee for Scientific Research and the British Council’s Young Scientist Programme). This research was funded through the 2018-2019 BiodivERsA joint call for research proposals, under the BiodivERsA3 ERA-Net COFOUND programme. MG was supported by the National Science Centre, Poland under BiodivERsA3 programme (2019/31/Z/NZ8/04028). MG thanks Alicja Rost and Ewa Zieliniewicz for their assistance in the laboratory.

## AUTHORSHIP

The study was conceived and designed by MG, DA and BB. Supervision of the long-term monitoring of bank vole populations in the region was by JMB and AB. Samples were collected in the field by JMB, AB, MA, MG, KT& JBB. The immunological analysis and laboratory work was conducted by DA, KT, JN, JMB, AB & MG. Data handling – MG. Statistical analysis was carried by JMB & MG. The manuscriptwas written by MG, BB, and JMB in consultation with all co-authors. MG, DA, AB & JMB revised the manuscript. All authors accepted the final manuscript version.

## AUTHOR BIO

Dr Grzybek is a parasitologist holding a position of associate professor at the Department of Tropical Parasitology, Institute of Maritime and Tropical Medicine, Medical University of Gdansk, Poland. His research interests include epidemiology and ecology of macro-and microparasites in rodents, especially in bank voles. He is also interested in host-parasite interactions and intrinsic and extrinsic factors that influence these relationships.

